# Causal inference gates corticostriatal learning

**DOI:** 10.1101/2020.11.05.369793

**Authors:** Hayley M. Dorfman, Momchil Tomov, Bernice Cheung, Dennis Clarke, Samuel J. Gershman, Brent L. Hughes

## Abstract

Attributing outcomes to your own actions or to external causes is essential for appropriately learning which actions lead to reward and which actions do not. Our previous work showed that this type of credit assignment is best explained by a Bayesian reinforcement learning model which posits that beliefs about the causal structure of the environment modulate reward prediction errors (RPEs) during action value updating. In this study, we investigated the neural circuits underlying reinforcement learning that are influenced by causal beliefs using functional magnetic resonance imaging (fMRI) while human participants (N = 31; 13 males, 18 females) completed a behavioral task that manipulated beliefs about causal structure. We found evidence that RPEs modulated by causal beliefs are represented in posterior putamen, while standard (unmodulated) RPEs are represented in ventral striatum. Further analyses revealed that beliefs about causal structure are represented in anterior insula and inferior frontal gyrus. Finally, structural equation modeling revealed effective connectivity from anterior insula to posterior putamen. Together, these results are consistent with a neural architecture in which causal beliefs in anterior insula are integrated with prediction error signals in posterior putamen to update action values.

**Significance Statement:** Learning which actions lead to reward – a process known as reinforcement learning – is essential for survival. Inferring the causes of observed outcomes – a process known as causal inference – is crucial for appropriately assigning credit to one’s own actions and restricting learning to effective action-outcome contingencies. Previous studies have linked reinforcement learning to the striatum and causal inference to prefrontal regions, yet how these neural processes interact to guide adaptive behavior remains poorly understood. Here, we found evidence that causal beliefs represented in the prefrontal cortex modulate action value updating in posterior striatum, separately from the unmodulated action value update in ventral striatum posited by standard reinforcement learning models.

## Introduction

We live in an uncertain environment where making flexible predictions about the occurrence of positive and negative events is necessary for maximizing rewards and minimizing punishments. Predictions are most accurate, and feedback most useful, when our own actions are responsible for the outcomes we receive. Thus, drawing inferences about the causes of outcomes is a critical component of credit assignment for learning.

We recently demonstrated that causal inferences can lead to asymmetric learning from positive and negative outcomes (Dorfman et al., 2019). Specifically, participants down-weighted outcomes when they could be attributed to the intervention of a latent agent. If the participants knew that the agent’s interventions tended to produce negative outcomes, then participants learned less from negative relative to positive outcomes. Conversely, if they knew that the agent’s interventions tended to produce positive outcomes, then participants learned less from positive relative to negative outcomes. These results demonstrate that people modulate the extent to which they learn depending on their beliefs about latent causes, and that these beliefs can be experimentally manipulated to produce learning biases.

The learning asymmetries reported in Dorfman et al. (2019) could be explained by a Bayesian model that assigns credit based on probabilistic inference over latent causes. Mechanistically, the model hypothesizes that reward prediction errors (RPEs) are weighted by the probability that the feedback was generated by the participant’s choice (rather than the latent agent intervention). The research reported here sought to test this mechanistic hypothesis using functional MRI.

Prior work in rodents, primates, and humans has shown that RPEs, which represent the difference between expected and received rewards, are encoded by dopaminergic neurons in the midbrain (Schultz et al., 1997), and prediction error information is projected to the striatum and frontal cortex (see Niv, 2009 for a review). In human neuroimaging, RPE signals are found in regions of the striatum, prefrontal cortex (PFC) and orbitofrontal cortex (OFC) (McClure et al., 2003; O’Doherty et al., 2003). Several lines of evidence suggest that striatal RPEs in particular may be sensitive to causal inference. For example, the caudate (dorsomedial striatum) and nucleus accumbens (ventral striatum) are sensitive to rewards that are chosen rather than passively received (Zink et al., 2004), and both the ventral and dorsal striatum are preferentially recruited during anticipation of a controllable (compared to an uncontrollable) outcome (Bhanji & Delgado; Leotti & Delgado, 2011; Stolz et al., 2020). Previous work has also shown diminished activation in the putamen (dorsolateral striatum) for trials where people believed they caused a loss (Späti et al., 2015).

To more directly investigate the interplay between causal inference and error-driven updating in the striatum, we measured brain activity in humans while they performed the task developed by Dorfman et al. (2019). We used a combination of model-based univariate analyses and structural equation modeling to map the information processing architecture posited by the Bayesian model.

## Materials and Methods

### Participants & Data Inclusion

Participants were recruited from the University of California at Riverside SONA study pool system. A total of 38 right-handed adults consented to the study, but seven of these individuals were not included due to unavoidable technical issues that resulted in missing and corrupted data, leaving 31 total participants for the current analyses (ages 18-24, mean = 19.4, 13 males, 18 females). Individual runs were evaluated for excessive head movement (> 4mm) over the duration of the run, but no runs were excluded for movement.

### Experimental Design and Statistical Analysis

The task was presented using PsychoPy software version 1.85.2 (Peirce, 2007), and displayed on a screen visible through a mirror attached to the head coil. Behavioral responses were collected with an MRI-compatible button box, and all participants used the index and middle finger of their right (dominant) hand to make responses. Prior to entering the scanner, participants received verbal instructions and completed a practice version of the task outside the scanner.

Participants completed a reinforcement learning task in which they encountered multiple learning environments. This task was modified from our previous behavioral task for use in the scanner (Fig. 1; Dorfman et al., 2019). Participants were instructed to imagine that they were mining for gold in the Wild West. On each trial, participants had to choose between one of two different colored mines using the button box. After making a decision (*choice period*: 2.5 s), an inter-stimulus fixation interval (*ISI*: jittered between 1.5 s-2.5 s) was displayed. Participants then viewed feedback of either a reward (gold) or loss (rocks) for 1.5 s. Each mine in a pair produced a reward with 70% or 30% probability. Participants were instructed that each reward yielded a small amount of bonus money and each loss resulted in subtraction of bonus money. In actuality, all participants received a $5 bonus at the end of the experiment.

**Figure 1.**
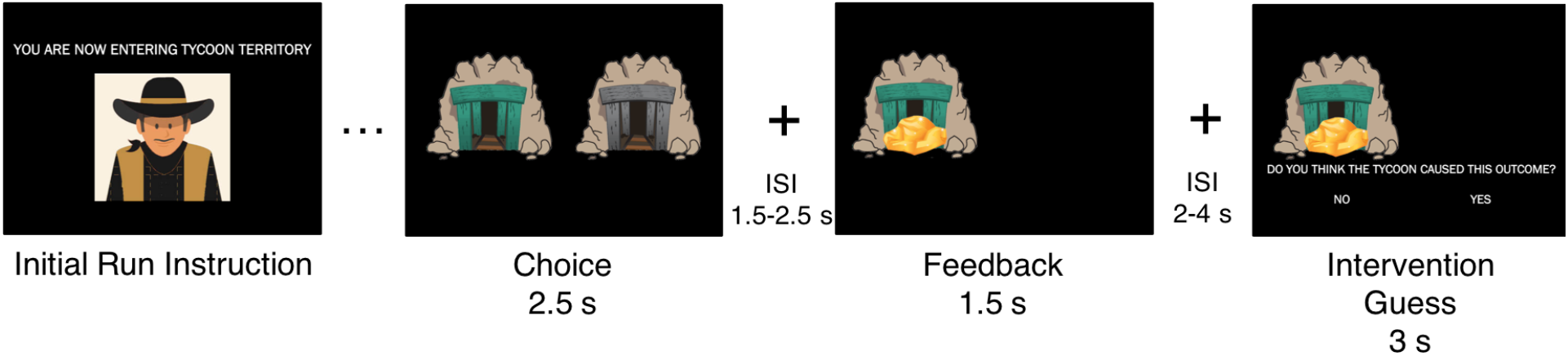
Task Schematic. At the start of each run, participants were told which territory they were in (Benevolent: “Tycoon” or Adversarial: “Bandit”). Trial components consist of a choice between two stimuli (2.5 s), a fixation inter-stimulus interval (ISI; 1.5-2.5 s), feedback (win: “Gold” or loss: “Rocks”; 1.5 s), a fixation ISI (2-4 s), and an intervention guess (3 s). At the end of each run, a fixation inter-run interval was presented for 10 s plus the residual amount of time from the stimulus choice and intervention guess events.

Participants completed four blocks of 30 trials (120 total trials) in different “mining territories.” A single block was presented for each functional run. Participants were instructed that different agents frequent each territory: a bandit will occasionally steal gold from the mines and replace it with rocks (adversarial condition) and a tycoon will occasionally leave extra gold in the mines (benevolent condition). During the task, participants completed two blocks of each condition, which were interleaved in a pseudorandomized fashion. The agents intervened on 30% of the trials, and participants were told this proportion explicitly at the start of the task, though they did not know unambiguously whether the agent intervened on any particular trial. While the underlying reward distributions (i.e., absent intervention) for the mines were 70% or 30%, the hidden agent intervened on 30% of trials (or 9 out of 30 trials). For example, the benevolent intervention produced rewards on 9 out of 30 trials and the adversarial intervention produced losses on 9 out of 30 trials, regardless of the participant’s choice. After feedback and the presentation of a second ISI (2-4 s), participants were asked whether they believed the outcome they received was a result of hidden agent intervention (binary response of “Yes” or “No”; 3s) and made their selection using the button box. The stimulus choice and intervention guess periods would end as soon as a button was pressed. Residual time from these periods was added to the subsequent inter-run interval (IRI; 10 s plus residual) at the end of the run/block.

### Computational Model

We used a computational model developed in our earlier work (Dorfman et al., 2019) to analyze the behavioral and neural data. The model proposes an update rule (Eq. 1), where *θ_t_* is the estimated value of a given action, *r_t_* is the reward outcome on each trial *t*, and *α_t_* is a parameter representing the learning rate that scales the reward prediction error (*r_t_ - θ_t_*):

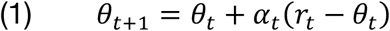

The learning rate is derived analytically to be consistent with Bayesian updating (see Dorfman et al., 2019, for more details). In particular, it depends on beliefs about the outcome’s two possible sources: the action’s intrinsic reward distribution and the latent agent’s intervention. The learning rate encodes the degree to which feedback should be attributed to the intrinsic reward distribution rather than to the latent agent, which is affected by the experimental condition and whether feedback was positive or negative. From now on, we will omit the trial subscript *t* to keep the notation uncluttered.

We use the indicator random variable *z* to denote whether the latent agent intervened (*z* = 1) or did not intervene (*z* = 0) on any given trial. In our experimental task, with probability *P*(*z* = 0), the decision maker receives a reward from the intrinsic distribution, *P*(*r*|*z* = 0), or with probability *P*(*z* = 1) they receive a reward from a distribution determined by the latent agent intervention, *P*(*r*|*z* = 1). A rational observer that is familiar with the task structure (as our participants are) can use this to infer the posterior probability that a latent agent did not intervene on a given trial, *P*(*z* = 0|r), which is given by Bayes’ rule:

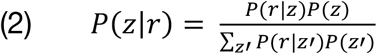

For simplicity, we summarize the posterior *P*(*z*|*r*) as a single quantity, *ψ* = *P*(*z* = 0|*r*), which reflects a trial-by-trial estimate of the decision maker’s belief that they caused the outcome, conditional on the past history of actions and rewards for the relevant task block. In Dorfman et al. (2019), we showed that the Bayes-optimal learning rate should be proportional to *ψ*. Intuitively, participants should be more sensitive to feedback when they believe it was drawn from the intrinsic reward distribution.

We fit three candidate models and compared them based on participant behavior (Table 1). In one model, we fixed the prior probability of hidden agent intervention at 30% to replicate the instructions that participants received in the task, while in another model, we set the prior probability of intervention as the mean of the participant’s subjective intervention judgments. The former we will refer to as the *fixed Bayesian model*, and the latter as the *empirical Bayesian model*. We hypothesized that either the fixed or empirical Bayesian models would best fit the behavior of our participants when compared to a four-learning rate model that fits separate learning rates for positive and negative prediction errors in each condition (e.g., positive Benevolent, negative Benevolent, positive Adversarial, and negative Adversarial).

**Table 1.**
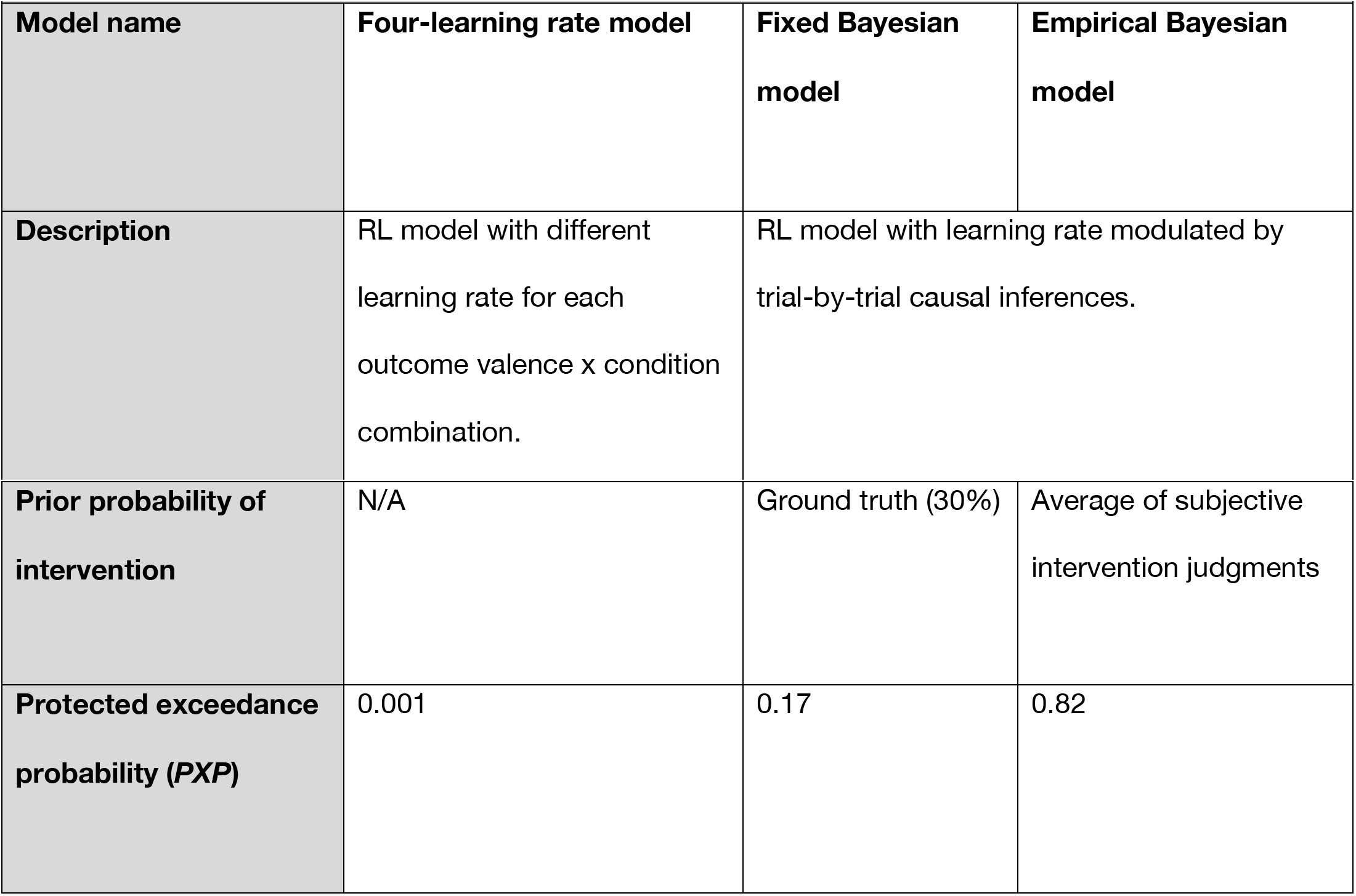
Computational model descriptions and model comparison results.

### Behavioral model comparison

We selected models for analyzing the neural data based on their fit to the behavioral data. We compared the empirical and fixed Bayesian models with the four-learning rate model using random-effects Bayesian model selection (Rigoux et al., 2014; Stephan et al., 2009). This procedure treats each participant as a random draw from a population-level distribution over models, which it estimates from the sample of model evidence values for each model. We approximated the log model evidence for each participant as LME = −0.5 * BIC, where BIC is the Bayesian information criterion based on maximum likelihood estimation of the free parameters of the given model. For our comparison metric, we report the *protected exceedance probability* (*PXP*), the probability that a particular model is more frequent in the population than all other models under consideration, taking into account the possibility that some differences in model evidence are due to chance.

### fMRI Data Acquisition & Preprocessing

Imaging data were collected on a 3.0 T Magnetom Prisma MRI scanner with the vendor 32-channel head coil (Siemens Healthcare, Erlangen, Germany) at the University of California Riverside Center for Advanced Neuroimaging. A T1-weighted high-resolution multi-echo magnetization-prepared rapid-acquisition gradient echo (ME-MPRAGE) anatomical scan (van der Kouwe et al., 2008) of the whole brain was acquired for each participant prior to any functional scanning (208 sagittal slices, voxel size = 0.8 x 0.8 x 0.8 mm, TR = 2400 ms, TE = 2.72 ms, TI = 1060 ms, flip angle = 8°, FOV = 256 mm). Functional images were acquired using a T2*-weighted echo-planar imaging (EPI) pulse sequence. In total, four functional runs were collected for each participant, with each run corresponding to a single task block, two for each condition (78 interleaved axial-oblique slices per whole brain volume, voxel size = 1.5 x 1.5 x 1.5 mm, TR = 2000 ms, TE = 32 ms, flip angle = 74°. Functional slices were oriented to a 30° tilt towards coronal from AC-PC alignment.

Functional data were preprocessed and analyzed using SPM12 (Wellcome Department of Imaging Neuroscience, London, UK). Each functional scan was realigned to correct for small movements between scans. The high-resolution T1-weighted ME-MPRAGE images were then co-registered to the mean realigned images and the gray matter was segmented and normalized to the gray matter of a standard Montreal Neurological Institute (MNI) reference brain (resampled voxel size 2 mm 343 isotropic). The functional images were then spatially smoothed with an 8 mm full-width at half-maximum (FWHM) Gaussian kernel, high-pass filtered at 1/128 Hz, and corrected for temporal autocorrelations using a first-order autoregressive model.

### fMRI Analysis

General linear models (GLMs) included impulse regressors that were convolved with the hemodynamic response function. Feedback onsets were modeled as the regressor of interest. All trial events besides the regressor of interest for a particular GLM were included as nuisance regressors, as well as motion estimates derived from the realignment procedure and run-specific intercepts. Trial-by-trial parameters from the computational models were included in relevant GLMs as parametric modulators, modeled at the time of feedback (see Table 2 for all GLM details).

**Table 2.** GLM Details. Regressor of interest is reported in regular text, nuisance regressors are italicized.

| Table 2: GLM Details |  |  |
| --- | --- | --- |
| Regressor name | Event | Which Trials |
| GLM 1: Reward prediction error (RPE) |  |  |
| RPE | Feedback Onset | Non-timeouts |
|  | *Feedback Onset* | Timeouts |
|  | *Trial Onset* | *All* |
|  | *Reaction Time / Trial Offset* | *All* |
|  | *Intervention Guess Onset* | *All* |
|  | *Intervention Guess Offset* | *All* |
| **GLM 2: Agency belief ($\psi$)** |  |  |
| $\psi$ | Feedback Onset | Non-timeouts |
|  | *Feedback Onset* | Timeouts |
|  | *Trial Onset* | *All* |
|  | *Reaction Time / Trial Offset* | *All* |
|  | *Intervention Guess Onset* | *All* |
|  | *Intervention Guess Offset* | *All* |
| **GLM 3: Agency-modulated reward prediction error ($RPE * \psi$)** |  |  |
| $RPE * \psi$ | Feedback Onset | Non-timeouts |
|  | *Feedback Onset* | Timeouts |
|  | *Trial Onset* | *All* |
|  | *Reaction Time / Trial Offset* | *All* |
|  | *Intervention Guess Onset* | *All* |
|  | *Intervention Guess Offset* | *All* |
| **GLM 4: RPE, agency belief, and agency-modulated RPE ($RPE, \psi, RPE * \psi$)** |  |  |
| $RPE, \psi, RPE * \psi$ | Feedback Onset | Non-timeouts |
|  | *Feedback Onset* | Timeouts |
|  | *Trial Onset* | *All* |
|  | *Reaction Time / Trial Offset* | *All* |
|  | *Intervention Guess Onset* | *All* |
|  | *Intervention Guess Offset* | *All* |

All group-level reported results are *t*-maps that have been whole-brain corrected at a voxel-wise threshold of *p* < 0.001 and cluster-corrected at *p* < 0.05, family-wise error (FWE). Region labels are based on the Harvard-Oxford Cortical and Subcortical Atlases and the SPM Automated Anatomical Labeling atlas (AAL2; Rolls et al., 2015; Tzourio-Mazoyer et al., 2002). Voxel-coordinates are reported in MNI space. ROIs for ψ from GLM 2 were defined as 4-mm spheres around the peak voxels in the corrected ψ contrast.

### Neural model comparison

Following our previous work, we compared GLMs using the same random-effects Bayesian model selection approach that we used for behavioral model comparison (Rigoux et al., 2014; Tomov et al., 2018). For voxel-wise Bayesian model selection, we computed log posterior odds maps, which are similar to posterior probability maps (Rosa et al., 2010). For each voxel, we performed random-effects Bayesian model selection using the BICs from that voxel alone, which gave us the (expected) posterior probabilities *r* of the two GLMs we compared, r_RPE_ (GLM 1: unmodulated RPE) and r_RPE*ψ_ = 1 - r_RPE_ (GLM 3: agency-modulated RPE). We then computed the log posterior odds for each voxel as log(r_RPE_ / r_RPE*ψ_). This produced a brain map with an interpretable scale: 0 means a voxel does not distinguish between the models, positive values mean the voxel favors unmodulated RPE representation (GLM 1), and negative values mean the voxel favors agency-modulated RPE representation (GLM 3). We thresholded the absolute values at 1.097, corresponding to a 0.75 or greater posterior probability favoring one model over the other. For subsequent analyses, ROIs for ventral striatum (VS) and posterior putamen (Put) were defined as unions of bilateral 4-mm spheres around the peak voxels in the log posterior odds map for the corresponding regions.

### Effective connectivity

For the update-specific effective connectivity analysis, we performed structural equation modeling (Ramsey et al., 2010; Spirtes, 2005). We extracted a beta series for feedback onset events using a GLM that was nearly identical to GLM 1, except that there was a separate feedback onset regressor on each (non-timeout) trial, and there was no parametric modulation. Using a beta series at feedback onset as opposed to the entire BOLD time course ensures that 1) we are only investigating functional coupling related to updating in response to feedback, and 2) our data points are relatively independent, which is a key assumption in structural equation modeling. Based on the univariate results reported below, we were interested in functional connectivity between four ROIs: the ψ ROIs in right inferior frontal gyrus (IFG) and right anterior insula (AI) from the ψ contrast in GLM 2, as well as the RPE ROI in bilateral ventral striatum (VS) and the RPE*ψ ROI in bilateral posterior putamen (Put) from the neural model comparison of GLMs 1 and 3. We also included three input variables: RPE, RPE*ψ, and ψ.

To explore the space of possible connectivity patterns, we used the IMaGES (independent multiple-sample greedy equivalence search) algorithm (Poldrack et al., 2011; Ramsey et al., 2010) from the TETRAD software package for causal modeling (Scheines et al., 1998). IMaGES is a modification of a greedy equivalence search (GES) algorithm (Meek, 1997) which starts with an empty causal graph and incrementally adds edges until it finds a graph that best fits the data according to the BIC. IMaGES adapts GES to modeling multiple datasets, such as multiple subjects in fMRI, by taking an average BIC across subjects (the *IMscore*).

We added the following required edges as prior knowledge for IMaGES: RPE → VS, RPE*ψ → Put, ψ → IFG, ψ → AI, since those ROIs were selected based on the corresponding input variables. This ensures that there is no circularity in the analysis (Kriegeskorte et al. 2009) and that any inferences about effective connectivity are not based on correlations between the variables used in the ROI selection procedure. We forbade the discovery of any edges between the input variables. We also restricted the search space to graphs that are compatible with our behavioral and computational modeling work. In particular, we only allowed edges between ROIs that represent variables which rely on one another for their computation: VS → Put, IFG → Put, Ins → Put, IFG → AI, AI → IFG.

We followed up with a confirmatory analysis using random-effects Bayesian model selection (Rigoux et al., 2014) on a restricted set of structural equation models that specifically test our hypotheses with respect to the winning causal graph identified by the IMaGES procedure. Fitting was performed using the **semopy** toolbox (Igolkina & Meshcheryakov, 2020) using Wishart maximum likelihood estimation, which assumes that the variables follow a multivariate normal distribution and their covariance follows a Wishart distribution. We then used the resulting BICs to perform random-effects Bayesian model selection as described previously. Note that this is a different kind of random-effects analysis than the one used by the IMaGES algorithm and therefore provides a complementary way of assessing and quantifying the result.

## Results

### Behavior & Computational Model

To verify that participants made attribution judgments consistent with the experimental manipulation, we examined hidden agent intervention judgments by outcome valence (win, loss) and condition (adversarial, benevolent). Replicating results from our previous work (Cohen et al., 2020; Dorfman et al., 2019), we found overall that participants were more likely to believe that negative outcomes were caused by the hidden agent compared to positive outcomes, *F*(1,30) = 12.65, p < 0.001. There was also a significant condition by outcome valence interaction, *F*(1,30) = 76.83, *p* < 0.0001, where participants were more likely to believe that the hidden agent had intervened after negative compared to positive outcomes in the adversarial condition, and after positive compared to negative outcomes in the benevolent condition.

We found that the empirical Bayesian model (*PXP* = 0.82) was superior to both the fixed Bayesian model (*PXP* = 0.17) and the four-learning rate model (*PXP* = 0.001). The model demonstrates our predicted two-way asymmetry between valence and condition driven by inferences about latent agent intervention, replicating our previous studies. We therefore used the empirical Bayesian model in all subsequent fMRI analyses.

To further interrogate our model, we also tested how the model predictions correspond to the subjective intervention judgments which were not used for fitting or model comparison. We found that the model-predicted learning rates were significantly lower for trials where participants believed that the hidden agent intervened, compared to trials where they believed that the hidden agent did not intervene, *t*(30) = −6.49, *p* < 0.0001, *d* = −1.17. In addition, participants’ judgments about intervention also showed a significant median point-biserial correlation with the intervention belief predicted by the model, *r* = 0.424, *p* < 0.0001.

All model-based fMRI analyses used trial-by-trial regressors extracted from the empirical Bayesian model (Table 1; Table 2).

### Reward Prediction Error Signals in Striatum and Medial Prefrontal Cortex

We first sought to verify that our task elicited the canonical striatal and prefrontal activation associated with RPEs. By entering a trial-by-trial RPE signal from the winning computational model (empirical Bayesian model) as a parametric modulator at feedback onset into a whole-brain GLM (GLM 1 in Table 2), we found robust activation in ventral striatum which extended to the ventromedial prefrontal cortex (Figure 2A, Table 3.1).

**Figure 2.**
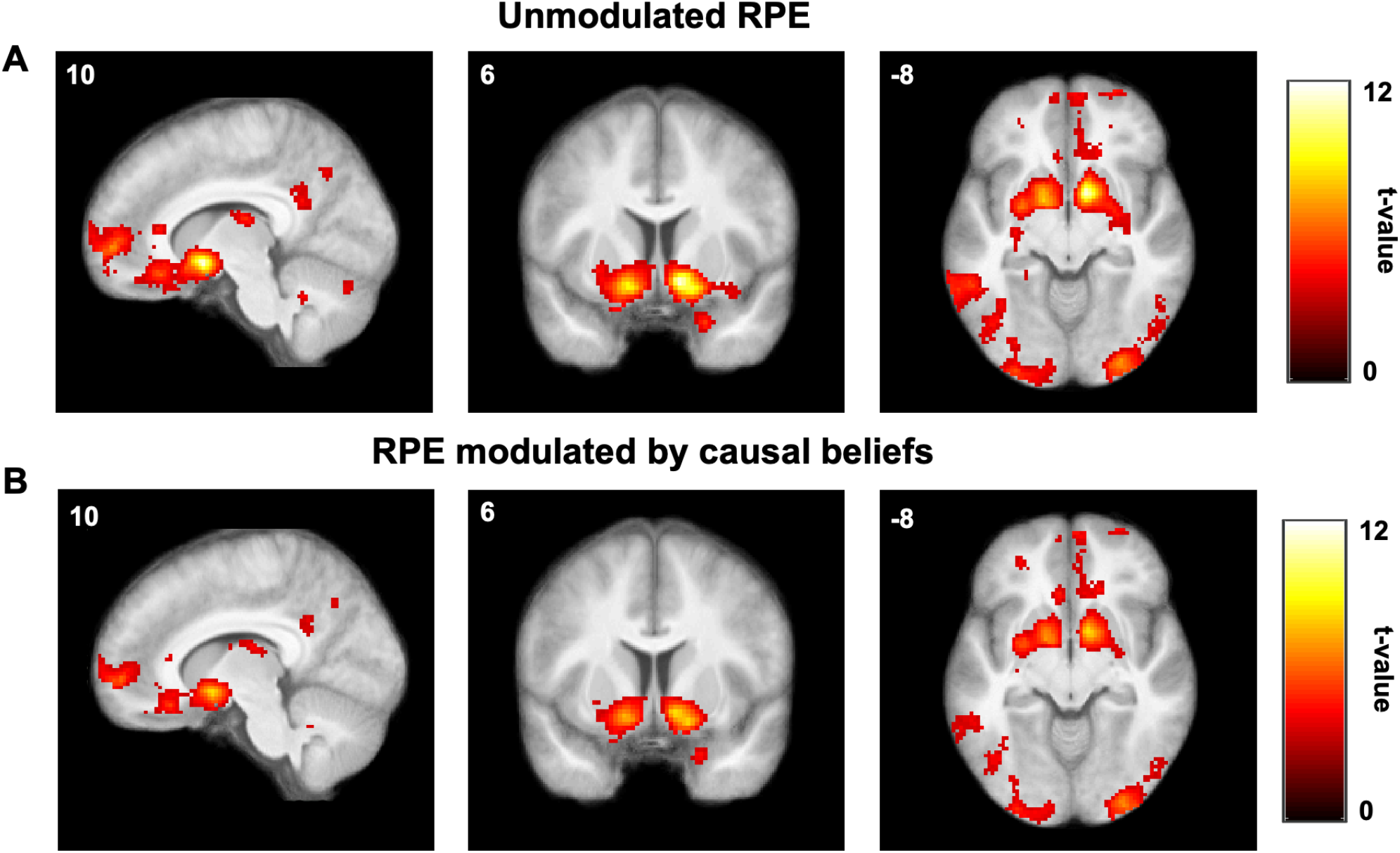
Reward prediction errors. Group-level statistical maps from GLM 1 (A) and GLM 3 (B) showing brain regions tracking unmodulated (A) and agency-modulated (B) reward prediction errors, respectively (single voxels thresholded at *p* < 0.001, whole-brain cluster FWE corrected at α = 0.05).

**Table 3.** Up to three subpeaks for the largest cluster are included.

| Table 3: BOLD activation |  |  |  |  |  |
| --- | --- | --- | --- | --- | --- |
| 3.1: Reward Prediction Error<br>(RPE) |  |  | MNI Coordinates |  |  |
| Region Label | Extent | t-value | x | y | z |
| Right Accumbens | 10142 | 11.405 | 10 | 6 | -10 |
| Left Cerebral Cortex | 10142 | 8.972 | -14 | 4 | -12 |
| Right Amygdala | 10142 | 7.477 | 24 | -6 | -14 |
| Left Cerebellum | 3592 | 6.357 | -38 | -74 | -34 |
| Left Cerebellum | 3592 | 6.166 | -26 | -48 | -26 |
| Left Inferior Temporal Gyrus | 3592 | 6.135 | -56 | -50 | -16 |
| Right Superior Occipital Gyrus | 2518 | 5.794 | 22 | -78 | 40 |
| Left Cuneus | 2518 | 5.396 | -12 | -78 | 40 |
| Left Precuneus | 2518 | 5.177 | -6 | -62 | 36 |
| Left Superior Frontal Gyrus | 732 | 5.583 | -18 | 24 | 54 |
| Left Superior Frontal Gyrus | 732 | 5.203 | -20 | 36 | 52 |
| Left Middle Frontal Gyrus | 732 | 4.693 | -32 | 24 | 58 |
| Right Superior Frontal Gyrus | 645 | 5.554 | 20 | 34 | 52 |
| Right Superior Frontal Gyrus | 645 | 4.863 | 28 | 44 | 46 |
| Right Superior Frontal Gyrus | 645 | 4.264 | 18 | 50 | 40 |
| Precentral Gyrus | 638 | 5.216 | 20 | -24 | 58 |
| Right Postcentral Gyrus | 638 | 4.720 | 24 | -46 | 68 |
| Postcentral Gyrus | 638 | 4.286 | 24 | -36 | 60 |
| Right Cerebellum | 168 | 4.909 | 42 | -72 | -36 |
| Right Cerebellum | 168 | 3.733 | 48 | -64 | -38 |
| Left Middle Frontal Gyrus | 560 | 4.831 | -48 | 48 | 6 |
| Left Middle Frontal Gyrus | 560 | 4.636 | -44 | 46 | 24 |
| Left Superior Frontal Gyrus | 560 | 4.625 | -12 | 68 | 10 |
| Left Precentral Gyrus | 527 | 4.752 | -26 | -30 | 66 |
| Left Precentral Gyrus | 527 | 4.491 | -22 | -22 | 72 |
| Precentral Gyrus | 527 | 4.458 | -14 | -30 | 62 |
| Left Angular Gyrus | 354 | 4.684 | -40 | -60 | 38 |
| Left Angular Gyrus | 354 | 4.447 | -44 | -70 | 30 |
| Left Angular Gyrus | 354 | 4.024 | -40 | -68 | 46 |
| Right Cerebellum | 203 | 4.608 | 18 | -68 | -28 |
| Cerebellar Vermis | 203 | 4.042 | 4 | -74 | -30 |
| Left Precuneus | 2518 | 3.563 | -8 | -70 | 38 |
| Right Rectal Gyrus | 10142 | 3.648 | 14 | 28 | -10 |
| Left Precentral Gyrus | 527 | 3.719 | -22 | -28 | 70 |
| 3.2: Belief about Agency (*psi*) |  |  | MNI Coordinates |  |  |
| Region Label | Extent | t-value | x | y | z |
| Right Inferior Frontal Gyrus<br>(pars triangularis) | 157 | 4.306 | 42 | 24 | 24 |
| Right Insula | 157 | 4.201 | 42 | 16 | 8 |
| 3.3: Agency-modulated RPE (*RPE\*psi*) |  |  | MNI Coordinates |  |  |
| Region Label | Extent | t-value | x | y | z |
| Right Caudate | 4766 | 8.204 | 10 | 6 | -8 |
| Left Putamen | 4766 | 7.407 | -14 | 6 | -10 |
| Right Hippocampus | 4766 | 6.899 | 24 | -8 | -14 |
| Right Inferior Occipital Gyrus | 802 | 6.685 | 30 | -92 | -8 |
| Right Inferior Occipital Gyrus | 802 | 5.268 | 50 | -80 | 0 |
| Right Inferior Occipital Gyrus | 802 | 4.489 | 38 | -88 | 0 |
| Left Cerebellum | 2197 | 6.405 | -36 | -76 | -34 |
| Left Cerebellum | 2197 | 5.937 | -26 | -46 | -26 |
| Left Inferior Occipital Gyrus | 2197 | 5.078 | -28 | -94 | -4 |
| Right Inferior Temporal Gyrus | 976 | 5.672 | 62 | -10 | -30 |
| Right Cerebellum | 976 | 5.165 | 36 | -40 | -26 |
| Right Inferior Temporal Gyrus | 976 | 5.110 | 62 | -30 | -16 |
| Left Putamen | 167 | 5.641 | -26 | -4 | 18 |
| Left Putamen | 167 | 4.351 | -30 | -4 | 8 |
| Right Superior Frontal Gyrus | 340 | 5.075 | 20 | 36 | 52 |
| Right Superior Frontal Gyrus | 340 | 4.531 | 28 | 44 | 46 |
| Right Superior Frontal Gyrus | 340 | 3.930 | 18 | 50 | 46 |
| Left Inferior Temporal Gyrus | 331 | 5.074 | -56 | -42 | -14 |
| Left Inferior Temporal Gyrus | 331 | 4.122 | -58 | -54 | -12 |
| Left Middle Temporal Gyrus | 331 | 3.642 | -64 | -46 | -4 |
| Right Thalamus | 275 | 5.032 | 14 | -16 | 18 |
| Right Caudate Nucleus | 275 | 4.731 | 18 | -4 | 24 |
| Right Caudate Nucleus | 275 | 3.498 | 10 | -4 | 16 |
| White Matter | 393 | 4.935 | 26 | -28 | 48 |
| White Matter | 393 | 4.770 | 20 | -28 | 58 |
| Right Postcentral Gyrus | 393 | 4.583 | 26 | -42 | 70 |
| Right Precuneus | 1048 | 4.808 | 4 | -62 | 42 |
| Left Cingulate | 1048 | 4.804 | -14 | -44 | 38 |
| Left Precuneus | 1048 | 4.766 | -8 | -58 | 36 |
| Left Superior Frontal Gyrus | 376 | 4.746 | -20 | 28 | 58 |
| Left Superior Frontal Gyrus | 376 | 4.394 | -16 | 50 | 46 |
| Left Middle Frontal Gyrus | 376 | 3.968 | -32 | 24 | 58 |
| Right Superior Occipital Gyrus | 171 | 4.621 | 22 | -78 | 40 |
| Right Superior Occipital Gyrus | 171 | 4.189 | 26 | -70 | 50 |
| Left Postcentral Gyrus | 247 | 4.222 | -22 | -34 | 68 |
| Left Precentral Gyrus | 247 | 4.207 | -28 | -24 | 58 |
| Left Paracentral Lobule | 247 | 4.170 | -16 | -20 | 70 |
| Left Fusiform Gyrus | 2197 | 3.528 | -44 | -66 | -10 |

Our computational model posits that the RPE is scaled by the agency belief, *ψ*, to obtain an agency-modulated RPE (RPE * *ψ*) which is used to update action values. Furthermore, previous studies in humans, nonhuman primates, and rodents have shown that different sub-regions of the striatum receive dopamine inputs encoding different kinds of RPE or value signals (Balleine et al., 2007; Matsumoto & Hikosaka, 2009; Menegas et al., 2015; Watabe-Uchida et al., 2012). This led us to hypothesize that different brain regions, and in particular different parts of the striatum, might be encoding the agency-modulated (RPE * *ψ*) and the unmodulated RPE. We therefore performed a similar whole-brain analysis with model-derived trial-by-trial estimates of RPE * *ψ* (GLM 3 in Table 2), which yielded a similar set of regions in ventral striatum and ventromedial prefrontal cortex (Figure 2B, Table 3.3).

To directly test the hypothesis that different parts of the striatum might be computing the unmodulated and agency-modulated RPE’s, we entered both regressors into a single GLM (GLM 4 in Table 2) and looked at the contrast between the two regressors (RPE - RPE * *ψ*). However, no voxels survived after correcting for multiple comparisons, likely due to the fact that the two regressors are highly correlated (average Pearson correlation of *p* = 0.81) and are trading off with each other (Mumford et al., 2015). While at first glance these results seem to suggest that the signals are not distinguishable using fMRI, the univariate analyses reported in this section may not be sufficiently sensitive to uncover a functional dissociation.

### Differential sensitivity of RPEs to causal beliefs in ventral striatum and posterior putamen

An alternative approach to testing whether a given brain region is sensitive to unmodulated versus agency-modulated RPEs is to perform model comparison between relevant GLMs (GLM 1: RPE and GLM 3: RPE * *ψ*). This approach has the advantage that it doesn’t make the implausible assumption that the same region might encode both forms of RPE to varying degrees; instead, it assumes that only one RPE is encoded by a region.

For each voxel in the striatum, we performed random-effects Bayesian model comparison (Rosa et al., 2010) and computed the log posterior odds as log(r_RPE_ / r_RPE*ψ_), where r_RPE_ is the posterior probability of GLM 1 (unmodulated RPE) and r_RPE*ψ_ is the posterior probability of GLM 3 (agency-modulated RPE). This resulted in a log posterior odds map quantifying the extent to which each voxel favors unmodulated RPE (GLM 1) over agency-modulated RPE (GLM 3) representation, with positive values favoring unmodulated RPE, negative values favoring agency-modulated RPE, and zero indicating the indifference point. We thresholded the map to only show voxels where the posterior probability of either model is 0.75 or greater. This revealed a graded pattern of RPE representation across striatum, with anterior ventral regions favoring unmodulated RPE representation (Figure 3A) and posterior dorsolateral regions favoring agency-modulated RPE representation (Figure 3B).

**Figure 3.**
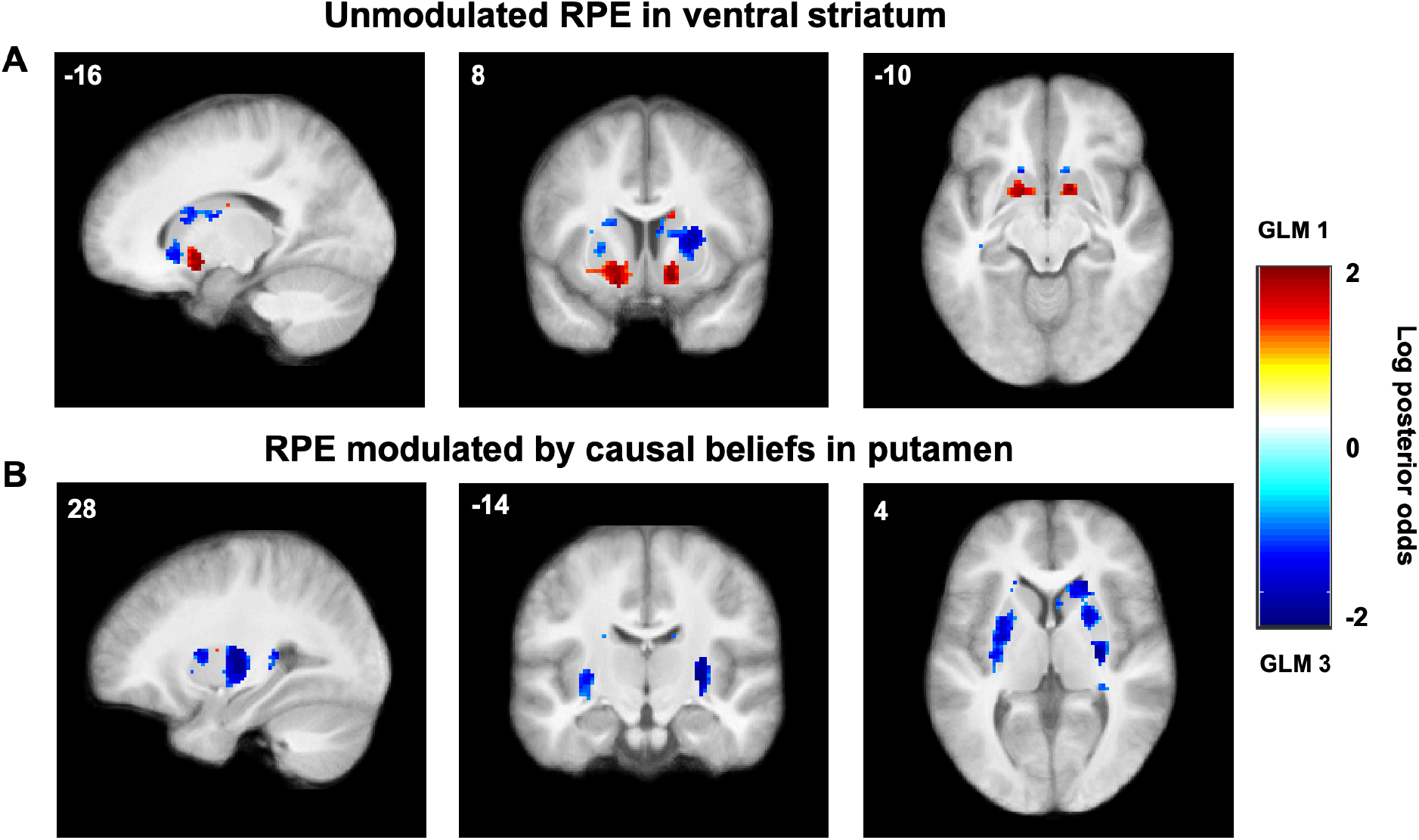
Log posterior odds of unmodulated versus agency-modulated RPEs. Log posterior odds map comparing GLM 1 (unmodulated RPE) and GLM 3 (agency-modulated RPE) showing striatal regions favoring (A) unmodulated RPEs in ventral striatum (green voxels) and (B) agency-modulated RPEs in posterior putamen (blue voxels). Single voxels thresholded at posterior probability 0.75 (log odds = 1.097) or greater. Color scales represent log posterior odds = log(r_RPE_ / r_RPE*ψ_), where r_RPE_ is the posterior probability of GLM 1 and r_RPE*ψ_ is the posterior probability of GLM 3. VS, ventral striatum (peaks [−16 8 −14] and [14 8 −12]); Put, posterior putamen (peaks [−28 2 2] and [28 −14 2]).

As an alternative way to quantify the same result, we performed model comparison in these regions of interest (ROIs). We defined a ventral striatum (nucleus accumbens) region as the union of two 4-mm spheres around the peak positive voxel (i.e. favoring unmodulated RPE), one in each hemisphere (MNI coordinates [−16 8 −14] and [14 8 −12]; we refer to this ROI as VS). We similarly defined a posterior putamen as the union of two 4-mm spheres around the peak negative voxel (i.e. favoring agency-modulated RPE), one in each hemisphere (MNI coordinates [−28 2 2] and [28 −14 2]; we refer to this ROI as Put). We then performed random effects Bayesian model selection comparing GLM 1 and GLM 3 in those two ROIs (note that this is not an independent confirmatory analysis, but rather a complementary way to quantify this same result which additionally takes into account the probability of the null hypothesis that there is no difference between GLM 1 and GLM 3). We found that ventral striatum favored unmodulated RPE representation (PXPs = 0.95 and 0.05 for GLM 1 and GLM 3, respectively) while posterior putamen favored agency-modulated RPE representation (*PXPs* = 0.13 and 0.87 for GLM 1 and GLM 3, respectively).

### Causal Beliefs in Anterior Insula and Inferior Frontal Gyrus

We next sought to identify regions that parametrically tracked *ψ*, the model quantity that tracks the degree to which participants believe they caused each outcome. To do so, we fitted a whole-brain GLM that included a parametric modulator for the belief about causal structure at feedback onset (*ψ*). Specifically, the quantity *ψ* represents (one minus) the posterior over hidden causes, with higher values for *ψ* accounting for a stronger sense of agency over the previous outcome. Using *ψ* as a single parametric modulator (GLM 2 in Table 2), a whole-brain analysis revealed a cluster in the right hemisphere with two subpeaks (Table 3.2): one in right anterior insula (Figure 4A; 157 voxels, peak [36 16 6]), and another in the triangular region of right inferior frontal gyrus (Figure 4B; 157 voxels, peak [42 24 24]). We defined two ROIs as 4-mm spheres around the corresponding peak voxels and henceforth refer to them as AI and IFG, respectively. These ROIs were then used for effective connectivity analyses, as discussed below.

**Figure 4.**
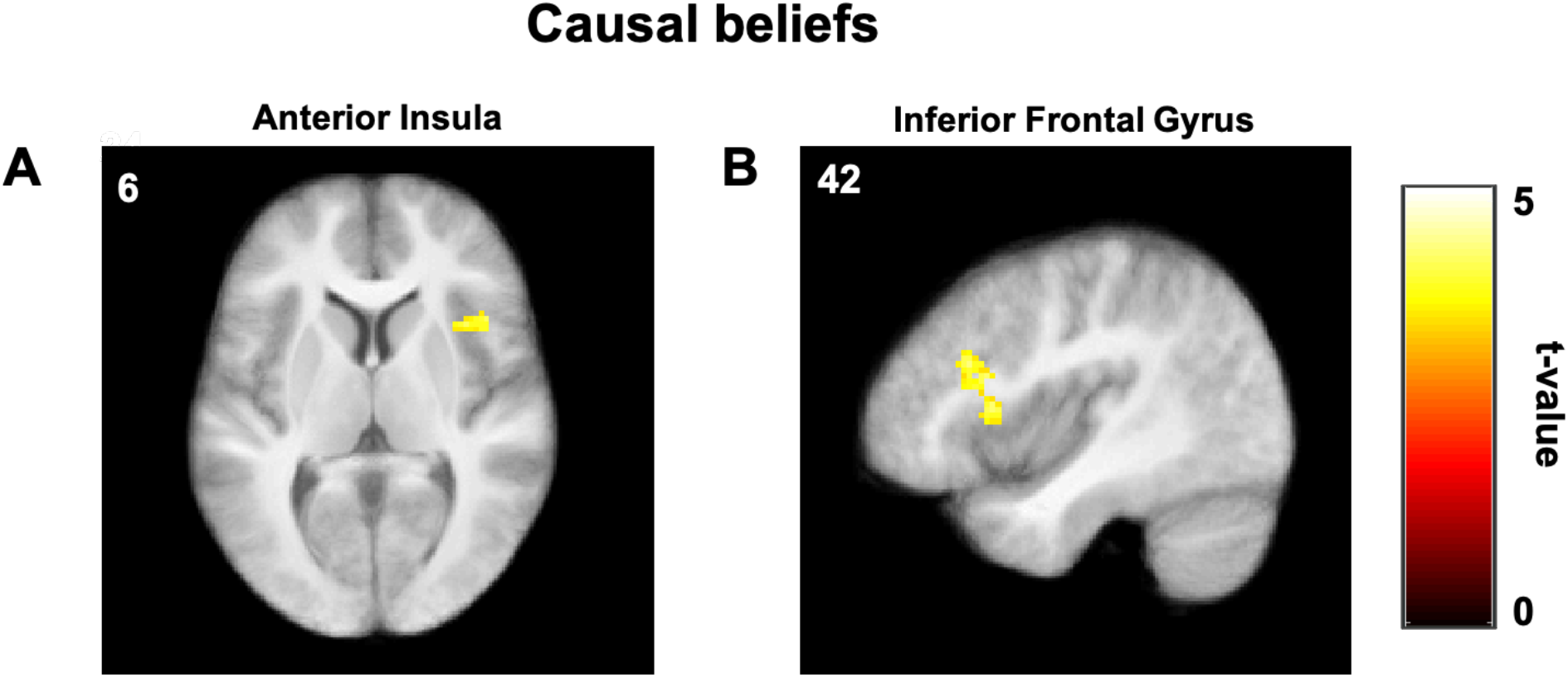
Beliefs about causal structure. Group-level statistical maps from GLM 2 showing brain regions tracking posterior beliefs about agency, *ψ*, single voxels thresholded at *p* < 0.001, whole-brain cluster FWE corrected at *α* = 0.05). Color scales represent t-values. (A) AI, anterior insula (peak [36 16 6]). (B) IFG, inferior frontal gyrus (peak [42 24 24]).

### Effective connectivity from anterior insula to posterior putamen but not from ventral stratum to posterior putamen during updating

Our computational model posits that causal beliefs *ψ*) and RPE are computed separately and then combined (RPE * *ψ*) to compute the action value update. Having identified regions that are associated with these quantities, we next sought to characterize the flow of information between those regions during updating. There are different possible patterns of connectivity between those regions that would be consistent with our model and with different hypotheses about corticostriatal architectures that have been put forward in the literature. In particular, actor-critic models of the basal ganglia favor parallel striatal architectures where state-value and action-value RPEs are computed independently in ventral striatum and dorsal striatum, respectively (Joel et al., 2002). This predicts that there will be no effective connectivity from ventral striatum to posterior putamen in our data. Conversely, some authors have reported evidence favoring a serial striatal architecture (Haber et al., 2000; Ikeda et al., 2013; Voorn et al., 2004) according to which results from computations in one part of the striatum are passed to another. This predicts that there will be effective connectivity from ventral stratum to posterior putamen, with posterior putamen integrating information about RPEs from ventral striatum. Similarly, posterior putamen might be integrating information about agency beliefs from either or none of the prefrontal regions associated with *ψ*.

To arbitrate between these possibilities, we performed effective connectivity analysis using structural equation modeling (Igolkina & Meshcheryakov, 2020; Ramsey et al., 2010; Spirtes, 2005) with a beta series extracted from feedback onset events. We searched the space of possible effective connectivity patterns using the IMaGES algorithm (see Methods) and found that in the pattern most consistent with the data, posterior putamen receives input from anterior insula, but not from ventral stratum nor inferior frontal gyrus (Figure 5A). Additionally, there is effective connectivity between anterior insular and inferior frontal gyrus, although the direction cannot be inferred from the data. Formally, there are two effective connectivity patterns (SEM 1A: IFG → AI; AI → Put; SEM 1B: AI → IFG, AI → Put) that correspond to causal graphs that are part of the same Markov equivalence class and hence cannot be disambiguated from our data. We confirmed this using Bayesian model selection (Rigoux et al., 2014) with the two equivalent SEMs (Figure 5B; PXPs = 0.57 and 0.43 for SEM 1A and SEM 1B, respectively).

**Figure 5.**
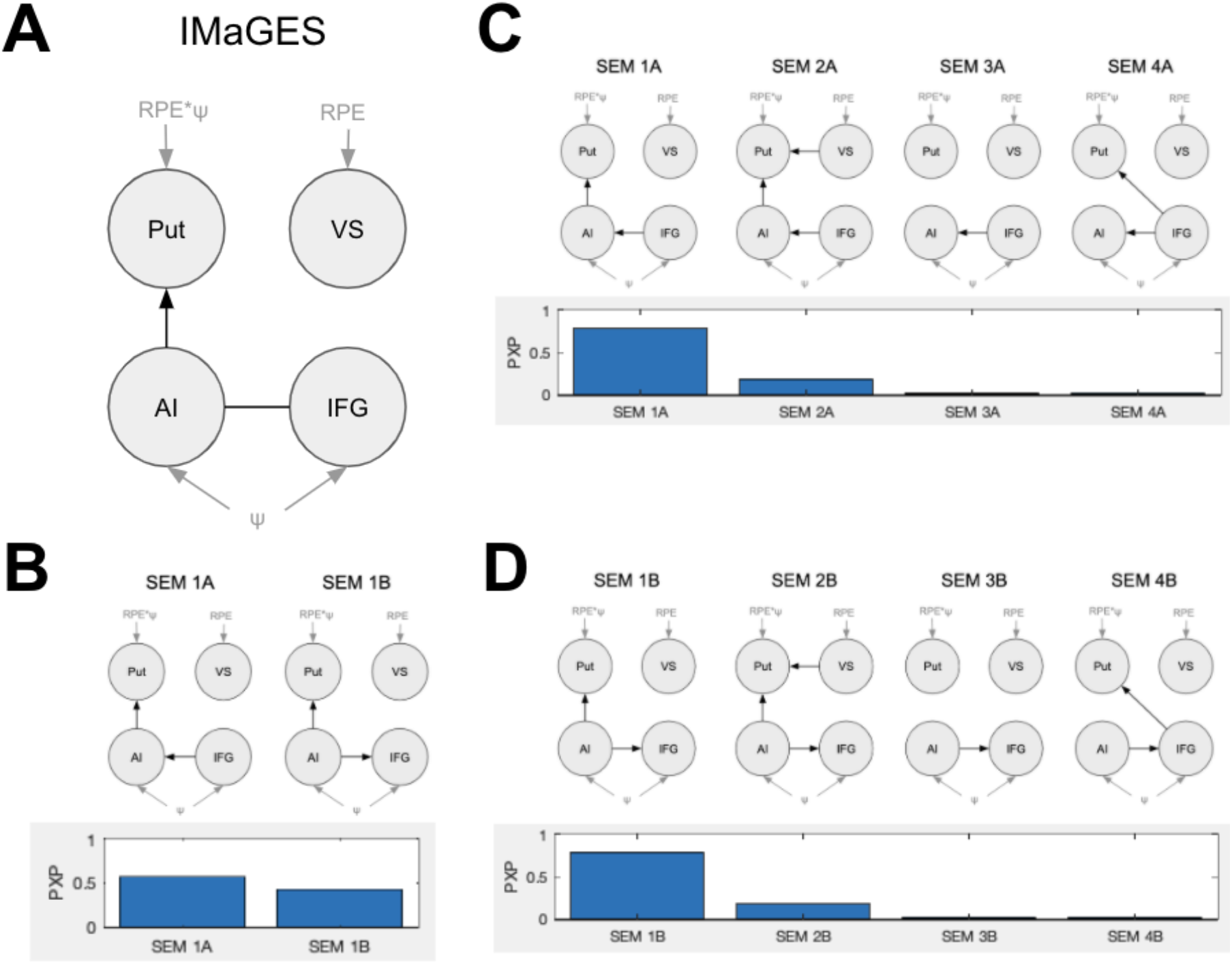
Effective connectivity analysis. (A) Winning connectivity pattern found by IMaGES algorithm using TETRAD. (B) Bayesian model selection with the two equivalent SEMs from A. (C and D) Bayesian model selection comparing modifications to SEM 1A (C) and SEM 1B (D). VS, ventral striatum; Put, posterior putamen; IFG, inferior frontal gyrus; AI, anterior insula. Gray nodes and edges indicate input variables. Undirected edges indicate that the either direction is consistent with the data. SEM, structural equation model. *PXP*, protected exceedance probability.

We performed additional model comparisons with modifications of the winning model to explicitly quantify the strength of the evidence favoring the above hypotheses (Figure 4C and 4D). In particular, we considered SEMs where posterior putamen receives input from ventral stratum (VS → Put), consistent with a serial striatal architecture (SEMs 2A and 2B). We also considered SEMs where posterior putamen receives no input from the cortical regions (SEMs 3A and 3B) or where it receives input from inferior frontal gyrus instead (SEMs 4A and 4B). Splitting these into two model comparisons to account for the ambiguity between SEMs 1A and 1B, we found that in both cases, the original winning model was favored (PXPs = 0.78, 0.18, 0.02, 0.02 for SEMs 1A, 2A, 3A, 4A and PXPs = 0.79, 0.18, 0.02, 0.02 for SEMs 1B, 2B, 3B, 4B, respectively). Our results thus point to a corticostriatal architecture for agency-modulated reinforcement learning where posterior putamen computes agency-modulated RPEs based on input about causal beliefs from anterior insula.

## Discussion

The present study used functional neuroimaging to uncover the neurobiological mechanisms that determine how causal beliefs modulate learning from positive and negative feedback and causal beliefs. To examine this, we measured differences in brain activation during a reinforcement learning task that manipulated participants’ causal beliefs about feedback. A whole-brain analysis revealed that anterior insula and inferior frontal gyrus represent causal beliefs, and that the striatum encodes both unmodulated and agency-modulated RPEs. However, a voxel-wise model comparison demonstrated that casual gating of RPEs follows an anatomical gradient, with ventral striatum representing unmodulated RPEs and posterior putamen representing agency-modulated RPEs. Finally, we analyzed alternative routes of how causal inference from cortical regions guides action selection in the striatum using structural equation modeling and found effective connectivity from anterior insula to posterior putamen but not from ventral striatum to posterior putamen, suggesting a possible corticostriatal circuit for agency-modulated reinforcement learning.

In addition, our results replicate our prior behavioral research and recapitulate canonical neural signatures of feedback learning. First, we successfully replicated our previous behavioral findings, showing increased learning for reward relative to loss outcomes in the adversarial condition and loss relative to reward outcomes in the benevolent condition, and that participant choice behavior was best explained by our hypothesized empirical Bayesian model. We also replicated our previous results which demonstrated that, collapsed across conditions, participants were more likely to believe that they caused positive outcomes and the latent agent caused negative outcomes, consistent with a self-serving bias (the attribution of good outcomes to oneself and bad outcomes to external forces; Campbell & Sedikides, 1999; Hughes & Zaki, 2015). Using the RPE from our winning model, we also found robust, canonical activation of the striatum and vmPFC at the time of feedback, demonstrating that participants are exhibiting prediction error activation consistent with the previous literature.

While these results reiterate that the striatum is integral for simple reinforcement learning, it was unclear whether additional regions contribute to a process of agency-modulated reinforcement learning. To investigate how causal beliefs influence feedback learning, we first conducted an exploratory whole-brain analysis using *ψ* as a parametric modulator to look for regions associated with this agency belief parameter. We found that activation in both the right anterior insula and right inferior frontal gyrus was associated with beliefs about self-agency.

A wealth of evidence implicates both the anterior insula and IFG in a variety of functions related to causal inference. In particular, the insula is recruited during agency judgments (David et al., 2008; Farrer & Frith, 2002; Sperduti et al., 2011), and both the insula and IFG are activated when rewards are personally chosen versus chosen by a computer (Romaniuk et al., 2019). There is also evidence that these regions are especially sensitive to negative self-attributions (Cabanis et al., 2013) and personally-undesirable estimation errors (Sharot et al., 2011) in particular. Relatedly, the AI and IFG have also been implicated in mismatch detection processes, with AI representing self-generated and externally-generated errors and IFG representing exclusively self-generated errors (Cracco et al., 2016). The IFG has also been shown to represent action-outcome likelihoods, and observed and executed goals or actions (Heiser et al., 2003; Iacoboni et al., 1999).

While AI and IFG seem to play a role in the representation of causal beliefs, our findings from the current study shed light on the specific computational processes associated with these regions during causal and latent state inference. Previously, we demonstrated that causal beliefs and their updating track with activation of the right AI and bilateral IFG, respectively, when participants made inferences about the causal relationships between cues, contexts, and outcomes (Tomov et al., 2018). In addition, representations of the belief about causal structure in this AI region tracked with variability in the sensitivity of subject behavior to the true causal structure of the environment (Tomov et al., 2018). Similarly, we previously reported an overlapping region of right AI that tracks likelihood estimates of group membership (Lau et al., 2020) Lau et al., (2020). Importantly, this estimate is calculated by gauging how similar the participant is to each of the possible group members, necessitating some form of self-representation. In addition, a recent study reports that right AI and IFG track causal structure in a self-concept network (Elder, Cheung, Davis, & Hughes, 2020) Together, these results suggest that AI and IFG perhaps represent inferences about the structure of the environment by calculating, integrating, and comparing information using the self as a reference point. Both the anterior insula and IFG in particular have been shown to be associated with a number of self-referential and self-related processes, including interoceptive awareness (Critchley et al., 2004; Mutschler et al., 2009), error awareness (Klein et al., 2013), mismatch detection, subjective confidence (Sherman et al., 2016), and self-awareness or consciousness (Braun et al., 2018; Craig, 2009; Karnath et al., 2005).

Our finding that dorsal and ventral subdivisions of the striatum encode different RPEs dovetails with a literature suggesting various action-related functional distinctions between these regions. Studies have overwhelmingly implicated the dorsal striatum in instrumental learning tasks, which involve outcome-linked, self-generated actions. For example, O’Doherty and colleagues (2004) reported ventral striatal activation for both Pavlovian and instrumental reward prediction errors (RPEs) but only dorsal striatal (anterior cingulate) activation for instrumental RPEs. Tricomi and colleagues (2004) showed that dorsal striatal activity was associated with a button press contingent with participants’ actions compared to button presses that were noncontingent, and increased activity in the dorsal striatum and OFC for actions that were more causal for outcomes. However, the dorsal striatum not only tracks the underlying reward-outcome contingencies, but there is also evidence that it tracks information about beliefs regarding these contingencies as well. For example, the dorsal striatum is activated for self-serving attributions, when participants believe they cause positive outcomes and someone or something else causes negative outcomes (Blackwood et al., 2003; Seidel et al., 2010).

Our results also provide an explanation for how causal beliefs in the insula integrate with agency-modulated RPEs in the dorsal striatum. Specifically, we found that anterior insula is functionally coupled to posterior putamen, which is consistent with anatomical studies showing white matter tracts between these regions (Ghaziri & Nguyen, 2018; Tian & Zalesky, 2018). This coupling suggests that beliefs about causal structure are passed from anterior insula to posterior putamen in order to compute the agency-modulated RPE that is ultimately used for updating action values. Additionally, this analysis showed that ventral striatum is not coupled to posterior putamen, which suggests that the two striatal regions receive RPE signals independently from each other (for example, from midbrain dopaminergic neurons), rather than ventral striatum passing the unmodulated RPE to posterior putamen. Thus our results are consistent with theories of parallel striatal architectures, which posit that ventral and dorsal striatum compute different types of RPEs in parallel (Joel et al., 2002) rather than serial striatal architectures, which posit that RPE information is passed from some parts of the striatum to others in a serial fashion (Haber et al., 2000; Ikeda et al., 2013; Voorn et al., 2004).

The current study also demonstrated functional connectivity between IFG and anterior insula, which is consistent with studies of white matter connectivity in humans (Cerliani et al., 2012; Deen et al., 2011). However, one limitation of our analysis method is that we cannot determine the connectivity direction between AI and IFG, so it remains unclear whether inferior frontal gyrus computes causal beliefs and passes them to anterior insula, or vice versa. This could be investigated in future studies using perturbation techniques such as TMS and tDCS. If the locus of the computation is in the inferior frontal gyrus, then perturbing both regions should have the same effect on behavior. Alternatively, if the locus of the computation is anterior insula, then only perturbing anterior insula should have an effect.

In summary, these results build on our previous behavioral and modeling work to shed light on the neural circuits that underpin agency-modulated reinforcement learning. Our results point to anterior insula and IFG as the origin regions of causal beliefs about agency, and posterior putamen as the region that integrates these beliefs with RPE signals. This agency-modulated RPE is distinct from, and computed in parallel with, the unmodulated RPE represented in ventral striatum. Together, these results bridge the gap between the rich literature grounding reinforcement learning in the basal ganglia circuitry and recent evidence of causal structure learning in the prefrontal cortex. By linking these regions in a way consistent with their prescribed combinational role, our findings pave the way to characterizing the neural circuits that allow humans to properly attribute outcomes to themselves or to external causes, and to use that knowledge for rational reward-based learning.

## Acknowledgements

Funding for this work was provided by the Office of Naval Research (N00014-17-1-2984 and N00014-17-1-2961), the Toyota Corporation, and the Alfred P. Sloan Foundation. The TETRAD software was developed by the Center for Causal Discovery at University of Pittsburgh, supported by grant U54HG008540.

